# Sequential Super-Resolution Imaging using DNA Strand Displacement

**DOI:** 10.1101/237560

**Authors:** Diane S. Lidke, Cheyenne Martin, Farzin Farzam, Jeremy S. Edwards, Sandeep Pallikkuth, Mathew R. Lakin, Keith A. Lidke

## Abstract

Sequential labeling and imaging in fluorescence microscopy allows the imaging of multiple structures in the same cell using a single fluorophore species. In super-resolution applications, the optimal dye suited to the method can be chosen, the optical setup can be simpler and there are no chromatic aberrations between images of different structures. We describe a method based on DNA strand displacement that can be used to quickly and easily perform the labeling and removal of the fluorophores during each sequence. Site-specific tags are conjugated with unique and orthogonal single stranded DNA. Labeling for a particular structure is achieved by hybridization of antibody-bound DNA with a complimentary dye-labeled strand. After imaging, the dye is removed using toehold-mediated strand displacement, in which an invader strand competes off the dye-labeled strand than can be subsequently washed away. Labeling and removal of each DNA-species requires only a few minutes. We demonstrate the concept using sequential dSTORM super-resolution for multiplex imaging of subcellular structures.

## Introduction

A key advantage in fluorescence microscopy is the ability to specifically label and image multiple targets within a cell. This is typically achieved through labeling of targets with spectrally distinct dyes with emissions that can be easily separated by filter-based detection. While this approach is achievable for many or the super-resolution imaging methods, the photo-physical requirements of the dye for each technique and the available dyes are not always ideally matched, placing limitations on multi-color super-resolution. For example, in single molecule localization microscopy (SMLM) it is desired to have low duty cycle and high number of photons per switching cycle. The best fluorophores for this are similar in spectra to Alexa647/Cy5 [1]. Additionally, the differential aberrations between spectral channels must be carefully accounted for in super-resolution imaging.

One approach for overcoming limitations in multi-color labeling is the use of sequential imaging, where the same fluorophore is used to image multiple structures in a label-image-remove process that is repeated for each target. This strategy has been demonstrated in several SMLM methods. Tam et al [2] used sequential labeling of antibodies and NaBH4 to quench remaining dyes while imaging using STORM [3]. Yi et al [4] used sequential antibody labeling and elution paired with dSTORM imaging [5]. Our own group demonstrated the use of sequential antibody labelling with bleaching and NaBH4 quenching for dSTORM imaging [6]. Each of these techniques requires a lengthy process between imaging steps to remove (photobleach or chemically quench) the fluorophore before labeling the next target. Another method, DNA-Exchange-PAINT [7] relies on transient binding of a dye-labeled imaging strand to a complementary docking strand to produce blinking. Sequential labeling can be done quickly by replacing the imaging strand in the buffer. However, DNA-PAINT suffers from high background due to the fluorescence imaging strand in the buffer, thus requiring TIRF or other selective illumination schemes. Also, the frame rate must be low, typically 100-300 ms, a factor approximately an order of magnitude lower than dSTORM imaging. An alternate strategy based on this approach uses semi-permanent binding of imaging strands to docking strands, allowing unbound imaging strands to be removed from the sample, thereby reducing background fluorescence [8]. A low salinity buffer with high concentrations of formamide is used to break hybridization for removal of the imaging strand.

Here we describe and evaluate a new sequential dSTORM labeling approach that allows for fast exchange of fluorescent labels while maintaining the inherent advantages of dSTORM - low background and higher data collection speeds. The concept relies on DNA strand displacement, which is a powerful method for designing enzyme-free reaction pathways to enable programmable, autonomous manipulation of DNA. Strand displacement has been used in a broad range of applications, including biological computing, molecular machinery, and in vivo biosensing [9–13]. Here we take advantage of the toehold-method of DNA strand displacement to sequentially add and remove the AF647 dye for multiplex dSTORM imaging without the need for enzymes or chemical treatments. We show that strand-displacement is an efficient and rapid means to exchange fluorescent dyes for sequential dSTORM imaging.

## Results

### DNA strand displacement for antibody labeling

Taking advantage of the properties of DNA interactions, we developed a labeling scheme for sequential dSTORM based on the method of toehold-mediated strand displacement. This is shown schematically in Figure 1. Using click chemistry, we directly conjugate the azide-modified “protector” strand to the secondary antibody. Upon addition of the complementary AF647-labeled “template” strand, the two DNA strands will hybridize, forming an AF647-labeled antibody. The “template” strand is removed by addition of the “invader” strand, which binds to the exposed toehold on the template strand and thereby nucleates a subsequent displacement reaction that displaces the template from the complex. In addition, the formation of additional base-pairs between the invader and the template strand provides a thermodynamic bias toward completion of the displacement process. Crucially, in our system this displacement reaction removes the AF647 label from the antibody as part of an inert waste complex that can then be washed away. After imaging and removal of the AF647 strand, the sample is relabeled with an orthogonal AF647-oligo that recognizes a different antibody via sequence-specific hybridization to its template strand, and the process can then be repeated. In this work, we describe two orthogonal DNA gate and invader strand sets that enable multiplex imaging. However, many more DNA sets could be constructed to allow for essentially any number of targets to be labeled.

**Figure 1:**
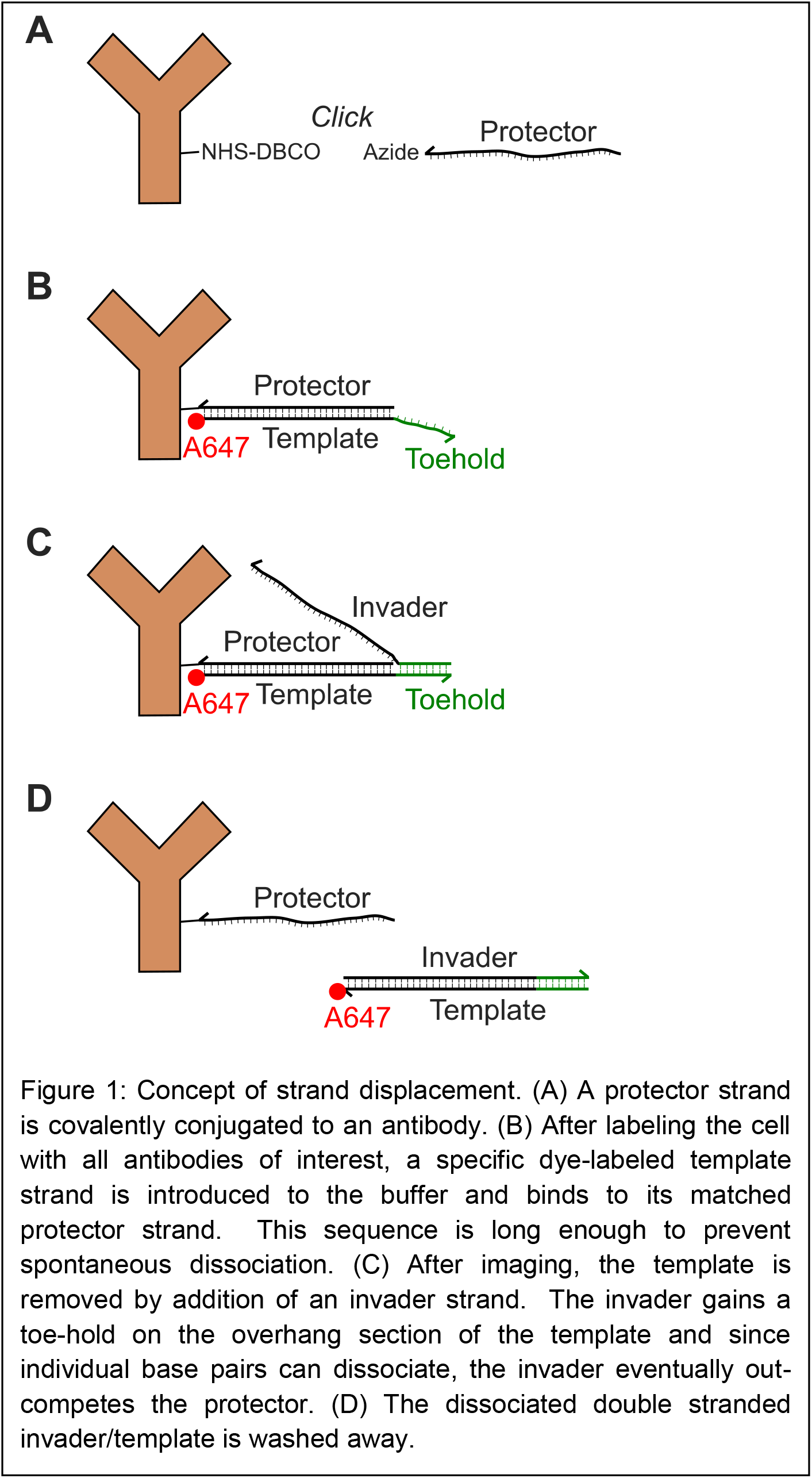
Concept of strand displacement. (A) A protector strand is covalently conjugated to an antibody. (B) After labeling the cell with all antibodies of interest, a specific dye-labeled template strand is introduced to the buffer and binds to its matched protector strand. This sequence is long enough to prevent spontaneous dissociation. (C) After imaging, the template is removed by addition of an invader strand. The invader gains a toe-hold on the overhang section of the template and since individual base pairs can dissociate, the invader eventually out-competes the protector. (D) The dissociated double stranded invader/template is washed away.

### AF647 is quickly and efficiently displaced by invader strand

The limiting step in sequential dSTORM is the potential for residual fluorescence from an initial target that could result in cross-talk artifacts for subsequent targets. We first established the ability of the invader strand to displace the AF647-template strand. HeLa cells were fixed and labeled with either anti-clathrin or anti-tubulin primary antibodies followed by the appropriate protector-labeled secondary antibody. Upon addition of AF647-template DNA, the oligo binds to the protector and robust labeling of the cells is achieved. To determine the kinetics of strand displacement, the cells are imaged over time during addition of the invader and the AF647 intensity for each cell was monitored (Fig 2, top). A rapid loss of AF647 is observed for both oligo sets with a greater than 95% reduction in intensity. To quantify the amount of residual cross-talk in sequential imaging, we compared dSTORM images of the same cell before (Fig. 2, middle) and after (Fig. 2, bottom) invader addition. Cross-talk, as determined by the total number of localizations, is found to be 2% and 5% for Set A and B, respectively.

**Figure 2:**
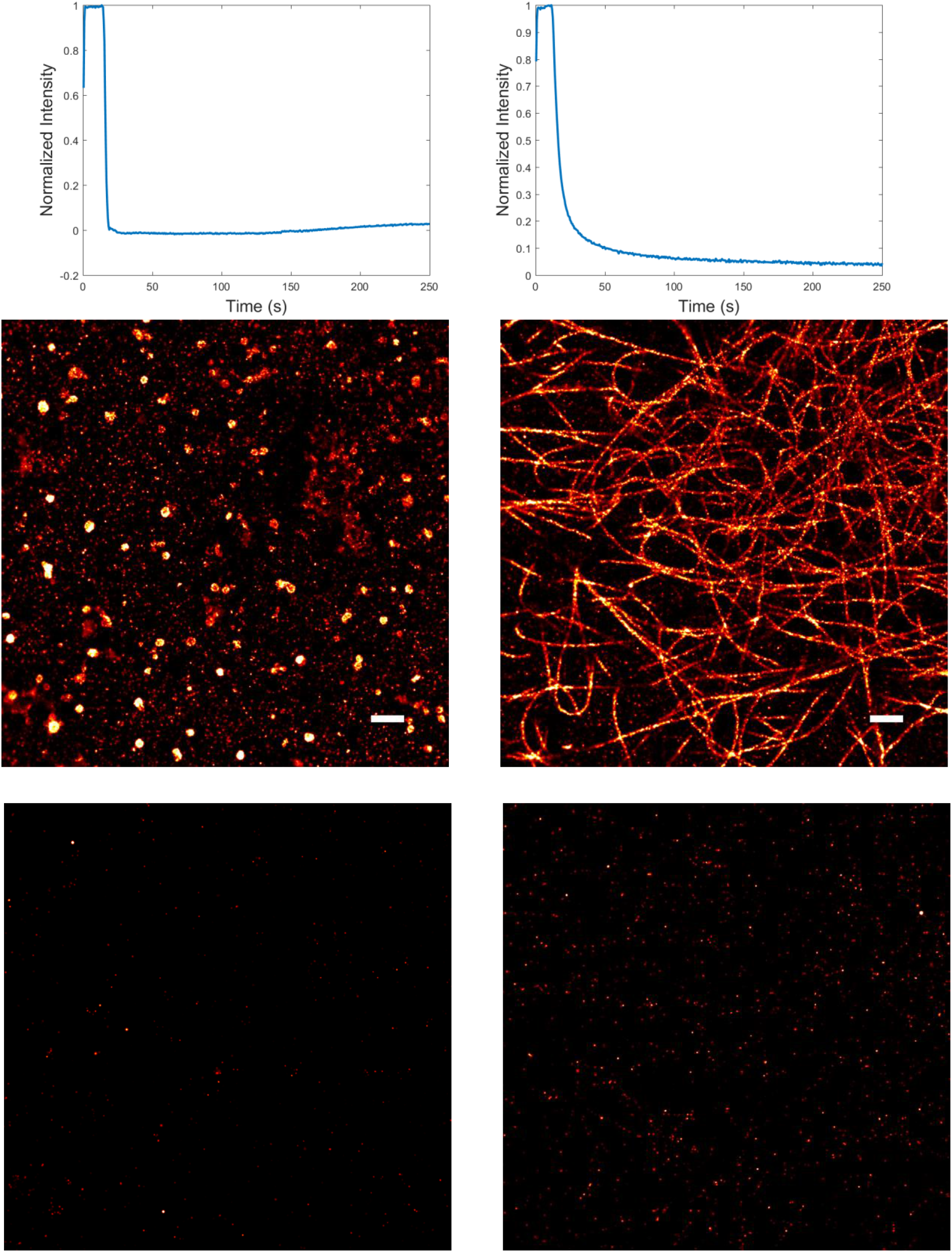
Invader Time Course and Residuals. (Left) Set A. (Right) Set B. (Top) Fluorescence intensity of a labeled cell sample as a function of time. The cell is imaged for ~ 20 s before adding invader. (Center) Super-resolution image before invader. (Bottom) Super-resolution image after invader. The residual fluorescence for Set A and Set B are 2% and 5% respectively.

### Sequential super-resolution imaging can be performed with DNA strand displacement

Since the two DNA sets are orthogonal (that is, none of the strands from set A will interact with strands from set B, and vice versa), they can be used to label distinct targets within the same cell. Figure 3 demonstrates the use of DNA strand displacement for sequential dSTORM. Cells were first labeled concurrently with clathrin and tubulin primary antibodies, followed by the appropriate protector-labeled secondary antibodies. The template for the first target is added and a super-resolution image acquired. Upon completion of the first round of imaging, the invader strand is added to remove the AF647-template from the first target and cells are washed. Next, the template strand to the second target is added and imaging of the second target is performed. We also performed the template labeling and imaging in the reverse order with similar results. A representative image from both labeling directions is shown in Figure 3.

**Figure 3:**
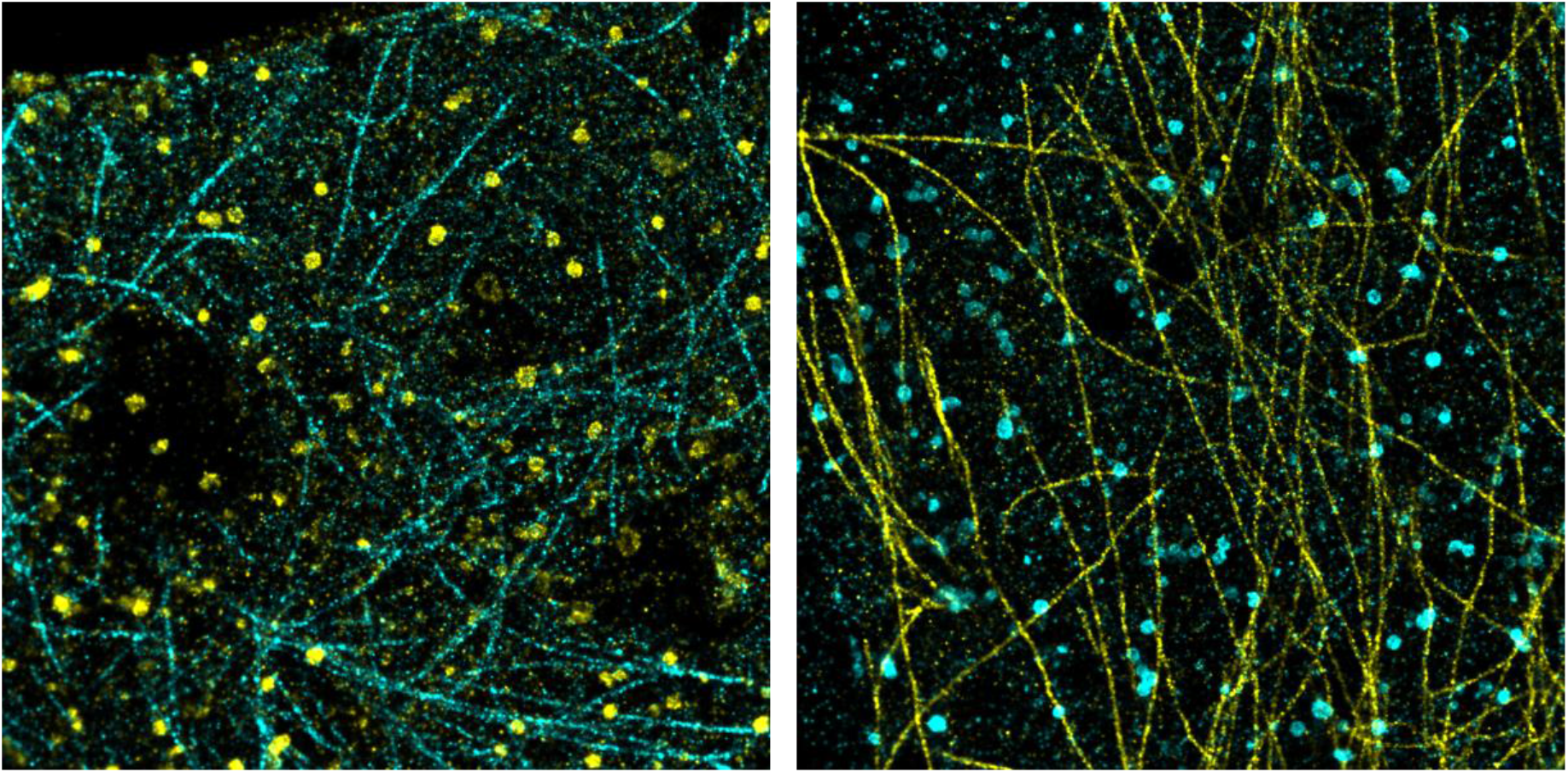
Sequential Imaging Results with Clathrin and Tubulin. Clathrin and tubulin are labeled and imaged using Set A and Set B, respectively. (Left) Set A (clathrin, yellow) is labeled and imaged and then removed with invader before labeling and imaging with Set B (tubulin, cyan). (Right) Set A and Set B labeling and imaging order is reversed (tubulin, yellow; clathrin, cyan).

### Discussion and Conclusion

We have shown the potential for DNA-strand displacement in providing a rapid and efficient labeling for multiplex sequential super-resolution. The exchange of labels requires about 10 minutes when performed by hand on the bench, although this could be reduced by approximately a factor of two using automated fluidics. This approach has the advantage over previous methods in that no enzymes, chemical treatments or photobleaching steps are required. The strand-displacement step is rapid thanks to the 8 nucleotide toehold, dramatically reducing the overall imaging time to the point where the total imaging time for dSTORM is dominated by the data collection. This is a significant improvement over our previous sequential dSTORM method [6] that required photobleaching, quenching by NaBH_4_, and an immunofluorescence labeling step between sequences, which resulted in approximately 2 hours for the complete removal and re-labeling steps per each structure.

The observed residual from strand displacement when applied to dSTORM imaging was as low as 2%. The residual cross-talk and speed of removal is similar to that of the denaturing approach used by Schueder *et al.* [8]. Whereas strand displacement requires an extra reagent, the invader stand, the denaturing approach uses a buffer with the odorous and suspected carcinogen, formamide. For applications that require even further reduction is cross-talk, strand displacement could be complemented by either photobleaching or dye quenching at the cost of a few minutes per cell for photobleaching or about 10 minutes per coverslip when quenching with NaBH_4_.

Since the DNA sets do not cross-react, sample preparation time is reduced by the ability to incubate all antibodies simultaneously and distinct targets are individually labeled at the time of imaging by addition of the corresponding AF647-template strand. While we have demonstrated this approach with two-target imaging, many more orthogonal DNA sets could be generated to increase multiplex capacity. Direct conjugation of the template strand to the primary antibody would allow for essentially limitless target combinations.

## Materials and Methods

### DNA Design

Four sets of non-interfering invader strand and gate sequences were designed using the NUPACK secondary structure prediction and design software [14–16] in conjunction with custom scripts to test and report any cross-reactivity between sets. Each strand displacement gate consists of a 38mer template strand and a 30mer protector strand. When hybridized this structure forms a 30 base pair duplex and an 8 nucleotide single-stranded toehold (using a long toehold promotes rapid nucleation of the strand displacement reaction). Each invader strand is a 38mer single strand that is complementary to the corresponding template strand. The super-resolution dye Alexa647 is attached to the 5’ end of each template strand, and the azide group for conjugation to the antibody is attached to the 3’ end of each protector strand. The two sets of nucleotide sequences, listed 5’ to 3’, are as follows:

Protector (This is bound to Antibody)

Template (This has Alexa647 for SR imaging)

Invader (Removes dye-labeled template)

Set A

GCCTGCTTTATCTCTGTTCTACTATTTCCG TT/3AzideN/

/5Alex647N/CGGAAATAGTAGAACAGAGATAAAGCAGGCAAACGAAA

TTTCGTTTGCCTGCTTTATCTCTGTTCTACTATTTCCG

Set B

GGGTCAAGTCAAAGTCAAGTATCAAGTCGG TT/3AzideN/

/5Alex647N/CCGACTTGATACTTGACTTTGACTTGACCCTTGATATT

AATATCAAGGGTCAAGTCAAAGTCAAGTATCAAGTCGG

### Materials

DNA oligonucleotides were purchased from Integrated DNA Technologies (Coralville, IA). Anti-alpha-Tubulin (Cat. T6074) primary antibody was purchased from Sigma Aldrich (St. Louis, MO). Anti-Clathrin (Cat. ab21679) antibody was purchased from Abcam (Cambridge, United Kingdom). Affini-Pure Donkey Anti-Mouse (DAMIG, Cat. 715-005-150) and Affini-Pure Donkey Anti-Rabbit (DARIG, Cat. 711-005-152) secondary antibodies were purchased from Jackson Immunoresearch (Westgrove, PA). 25mm #1.5 coverslips (Cat. CS-25R15) were from Warner Instruments (Hamden, CT). Bovine Serum Albumin and Triton X-100 were purchased from Sigma Aldrich (St. Louis, MO). Paraformaldehyde and glutaraldehyde were obtained from Electron Microscopy Sciences (Hatfield, PA). Image-iT™ FX Signal Enhancer (Cat. I36933) was purchased from Life Technologies (Carlsbad, CA).

### Antibody-DNA Conjugation

“Protector” DNA strands were coupled to antibodies through a two-step process. First, antibodies (1 mg/mL) were reacted with dibenzocyclooctyne (DBCO)-*sulfo*-NHS Ester at a 6:1 ratio in PBS + 100 mM NaHCO_3_ for 30 min at room temperature with gentle mixing. Unreacted DBCO-sulfo-NHS ester was removed using a Pro-Spin Column (Cat. CS-800, Princeton Separations) according to manufacturer’s instructions. This resulted in ~3:1 DBCO:Ab final ratio. Second, 3’-azide-modified DNA is covalently bound to the DBCO group via a Cu-free click reaction. DNA-azide was added to Antibody-DBCO at a 4:1 ratio and allowed to incubate overnight at 4°C. The next day, the sample volume was adjusted to 500 μl with PBS and unreacted DNA was removed using an Amicon Microcentrifuge Filter (100 kDa, Cat. UFC510024, Millipore) according to manufacturer’s instructions. This resulted in a final Protector:Ab ratio of ~1.5-2. The final sample was stored at 4°C for up to one month.

### Cell Fixation and Labeling

HeLa cells were cultured in Dulbecco’s Modified Eagle Medium (DMEM) with 10% fetal calf serum and pen/strep. For imaging, cell were cultured on piranha-cleaned 25 mm coverslips within a 6-well plate at 200,000 cells per well. Cell were allowed to adhere to the glass for 12-16 hr before fixation as described in Valley et al [6]. Briefly, cells were fixed with 0.6% Paraformaldehyde-0.1% Glutaraldehyde-0.25% Triton X-100 for 1 min and immediately followed with 4% Paraformaldehyde-0.2% Glutaraldehyde for 2 hr. Cells were then incubated with 0.1% NaBH_4_ for 5 min, washed with 10 mM Tris in PBS, and blocked with 4% BSA-0.1% Triton X-100 in PBS for 1 hr. Finally, cells were incubated with Signal Enhancer for 15 min. For labeling, antibodies were diluted in 2% BSA-0.05% Triton X-100 in PBS. Cells were incubated with primary antibodies at 10 μg/mL for 1 hr, followed by extensive washes then incubation with appropriate oligo-conjugated secondary antibodies at 5 μg/mL for 1 hr. Cells were again washed and post-fixed with 2% PFA for 15 min. Final washes of 10 mM Tris-PBS and then PBS were performed and coverslips were stored in PBS at 4°C until imaging. Fixation and antibody labeling was performed at RT.

### Adding and removing templates

Each coverslip was mounted on an Attofluor cell chamber (A-7816, life technologies) and washed four times with 1 mL of PBS. To label target structure with Alexa647, 400 μL of template solution (4.87 μM template in PBS) was added to the coverslip for 5 min. Excess template was removed by four washes with 1 mL of PBS and the coverslip was covered with 1.2 mL of dSTORM imaging buffer. Imaging buffer consisted of an enzymatic oxygen scavenging system and primary thiol: 50 mM Tris, 10 mM NaCl, 10% w/v glucose, 168.8 U/ml glucose oxidase (Sigma #G2133),1404 U/ml catalase (Sigma #C9332), and 32 mM 2-aminoethanethiol (MEA), pH 8. The chamber was sealed by placing an additional coverslip over the top.

To remove template after imaging a specific structure, the cell chamber was removed from the microscope and the coverslip was washed four times with 1 mL of PBS, followed by incubation with 4.87 μM invader in 400 μL of PBS to remove the template. To image the next structure, cells were washed four times with PBS, followed by addition of the template specific to the new target for 5 min, four washes with PBS and final addition of imaging buffer.

### Microscopy and Optical Setup

The imaging system is custom-built from an inverted microscope (IX71, Olympus America Inc.). A *xyz* piezo stage (Mad City Labs, Nano-LPS100) mounted on a *x-y* manual stage is installed on the microscope for cell locating and brightfield registration. The trans-illumination halogen lamp equipped with the microscope is used for collecting the brightfield images. The trans-illumination lamp power supply is connected to an analog/digital I/O board and controlled by a NI-DAQ card that allows the computer to adjust the lamp intensity by sending an analog output voltage from 0 V to 5 V to the lamp power supply. A 642 nm laser (collimated from a laser diode, HL6366DG, Thorlabs) is coupled into a multi-mode fiber (P1-488PM-FC-2, Thorlabs) and focused onto the back focal plane of the objective lens with 1.45 NA (UAPON 150XOTIRF, Olympus America Inc.) for data collection. A quad-band dichroic and emission filter set (LF405/488/561/635-A; Semrock, Rochester, NY) was used for sample illumination and emission. Emission light was filtered using a band-pass filter (685/45, Brightline) and collected on an iXon 897 electron-multiplying charge-coupled device (EM CCD) camera (Andor Technologies, South Windsor, CT). The EMCCD gain was set to 100, and frames were 256 × 256 pixels (for each channel) with a pixel size of 0.1067 μm. The emission path includes a quad band optical filter (Photometrics, QV2-SQ) with 4 filter sets (600/37, 525/45, 685/40, 445/45, Brightline) and the EM CCD camera mentioned above. All of the instruments are controlled by custom-written software in MATLAB (MathWorks Inc.).

### Super-resolution Imaging

When imaging each label, a brightfield reference image was acquired in addition to the position of each cell imaged. During data acquisition, the 642 nm laser was used at ~ 1 kW/cm^2^ to take 20 sets of 2,000 frames at 60 Hz. After imaging all target cells, the coverslip chamber was removed from the microscope and exchange of label was performed as described above. After labeling, the coverslip holder was placed back on the microscope and each cell was located and re-aligned using the saved brightfield reference image as described in [6]. Imaging was performed as described in *Microscopy and Optical Setup.* Data was analyzed via a 2D localization algorithm based on maximum likelihood estimation (MLE) [17]. The localized emitters were filtered through thresholds of a maximum background photon counts at 200, a minimum photon counts per frame per emitter at 250, and a data-model hypothesis test [18] with a cutoff p-value at 0.01. The accepted emitters were used to reconstruct the SR image. Each emitter was represented by a 2D-Gaussian with *σ_x_* and *σ_y_* equal to the localization precisions, which were calculated from the Cramér-Rao Lower Bound.

### Measurement of invader time course

Cells were labeled and template bound to the protector as described above. Imaging was performed in PBS buffer to match the typical conditions used for strand displacement. Data was collected at 2 Hz using low laser intensity to prevent photobleaching. Images were collected for ~ 20 s to establish a photobleaching rate and then invader strand was added to a final concentration of 4.87 μM during the time series acquisition. Time course plots were generated from the raw image data by subtracting from each frame a scalar background offset using the DipImage [19] function ‘backgroundoffset’. The sum intensity of each frame was calculated and normalized to the highest value in the intensity trace.

### Calculation of cross talk

Super-resolution imaging was performed on template-labeled cells, before and after treatment with invader. The measured data was then analyzed to determine the number of localizations. The ratio of number of localizations after treatment with invader to that before invader was determined as a measure of crosstalk. A crosstalk in the range of 2% to 5% was detected for the experiments reported here.

## Acknowledgments

We thank Marjolein Meddens for many helpful discussions and Mark Olah for connecting us to the molecular computing group at UNM. This work was supported by NIH 1R21EB019589, the New Mexico Spatiotemporal Modeling Center (NIH P50GM085273). M.R.L was additionally supported by the National Science Foundation grants 1525553, 1518861, and 1318833. We thank Shayna Lucero for assistance with cell culture. We gratefully acknowledge use of the University of New Mexico Comprehensive Cancer Center fluorescence microscopy core, as well as the NIH P30CA118100 support for these cores.

## References

1 Dempsey, G.T., J.C. Vaughan, K.H. Chen, M. Bates, and X.W. Zhuang, Evaluation of fluorophores for optimal performance in localization-based super-resolution imaging. Nature Methods, 2011. 8(12): p. 1027-+.

2 Tam, J., G.A. Cordier, J.S. Borbely, Á. Sandoval Álvarez, and M. Lakadamyali, Cross-Talk-Free Multi-Color STORM Imaging Using a Single Fluorophore. PLoS ONE, 2014. 9(7): p. e101772.

3 Rust, M.J., M. Bates, and X.W. Zhuang, Sub-diffraction-limit imaging by stochastic optical reconstruction microscopy (STORM). Nature Methods, 2006. 3(10): p. 793–795.

4 Yi, J., A. Manna, V.A. Barr, J. Hong, K.C. Neuman, and L.E. Samelson, madSTORM: a superresolution technique for large-scale multiplexing at single-molecule accuracy. Molecular Biology of the Cell, 2016. 27(22): p. 3591–3600.

5 Heilemann, M., S. van de Linde, M. Schuttpelz, R. Kasper, B. Seefeldt, A. Mukherjee, P. Tinnefeld, and M. Sauer, Subdiffraction-resolution fluorescence imaging with conventional fluorescent probes. Angewandte Chemie-International Edition, 2008. 47(33): p. 6172–6176.

6 Valley, C.C., S. Liu, D.S. Lidke, and K.A. Lidke, Sequential Superresolution Imaging of Multiple Targets Using a Single Fluorophore. Plos One, 2015. 10(4).

7 Jungmann, R., M.S. Avendano, J.B. Woehrstein, M.J. Dai, W.M. Shih, and P. Yin, Multiplexed 3D cellular super-resolution imaging with DNA-PAINT and Exchange-PAINT. Nature Methods, 2014. 11(3): p. 313–U292.

8 Schueder, F., M.T. Strauss, D. Hoerl, J. Schnitzbauer, T. Schlichthaerle, S. Strauss, P. Yin, H. Harz, H. Leonhardt, and R. Jungmann, Universal Super-Resolution Multiplexing by DNA Exchange. Angewandte Chemie International Edition, 2017. 56(14): p. 4052–4055.

9 Groves, B., Y.-J. Chen, C. Zurla, S. Pochekailov, J.L. Kirschman, P.J. Santangelo, and G. Seelig, Computing in mammalian cells with nucleic acid strand exchange. Nature Nanotechnology, 2015. 11: p. 287.

10 Chen, Y.-J., N. Dalchau, N. Srinivas, A. Phillips, L. Cardelli, D. Soloveichik, and G. Seelig, Programmable chemical controllers made from DNA. Nature Nanotechnology, 2013. 8: p. 755.

11 Qian, L. and E. Winfree, Scaling Up Digital Circuit Computation with DNA Strand Displacement Cascades. Science, 2011. 332(6034): p. 1196–1201.

12 Zhang, D.Y. and G. Seelig, Dynamic DNA nanotechnology using strand-displacement reactions. Nature Chemistry, 2011. 3: p. 103.

13 Yurke, B., A.J. Turberfield, A.P. Mills Jr, F.C. Simmel, and J.L. Neumann, A DNA-fuelled molecular machine made of DNA. Nature, 2000. 406: p. 605.

14 Wolfe, B.R., N.J. Porubsky, J.N. Zadeh, R.M. Dirks, and N.A. Pierce, Constrained Multistate Sequence Design for Nucleic Acid Reaction Pathway Engineering. Journal of the American Chemical Society, 2017. 139(8): p. 3134–3144.

15 Wolfe, B.R. and N.A. Pierce, Sequence Design for a Test Tube of Interacting Nucleic Acid Strands. ACS Synthetic Biology, 2015. 4(10): p. 1086–1100.

16 Zadeh, J.N., C.D. Steenberg, J.S. Bois, B.R. Wolfe, M.B. Pierce, A.R. Khan, R.M. Dirks, and N.A. Pierce, NUPACK: Analysis and design of nucleic acid systems. Journal of Computational Chemistry, 2011. 32(1): p. 170–173.

17 Smith, C.S., N. Joseph, B. Rieger, and K.A. Lidke, Fast, single-molecule localization that achieves theoretically minimum uncertainty. Nature Methods, 2010. 7(5): p. 373–U52.

18 Huang, F., S.L. Schwartz, J.M. Byars, and K.A. Lidke, Simultaneous multiple-emitter fitting for single molecule super-resolution imaging. Biomedical Optics Express, 2011. 2(5): p. 1377–1393.

19 Hendriks, C.L., L. Van Vliet, B. Rieger, G. van Kempen, and M. van Ginkel, DIPimage: a scientific image processing toolbox for MATLAB. Quantitative Imaging Group, Faculty of Applied Sciences, Delft University of Technology, Delft, The Netherlands, 1999.

